# A simple and efficient CTAB plate-based protocol for genomic DNA extraction from crop plants

**DOI:** 10.64898/2026.01.06.697759

**Authors:** Ha Quach, Gabriel de Bernardeaux, Duc Nguyen, Seema Sahay, Khang Hoang

## Abstract

Modern plant breeding and molecular genetics rely on genotyping large populations for applications such as genome editing and genomic selection. However, current DNA extraction methods often require expensive equipment or significant manual labor, which limits their scalability. The objective of this study was to develop a scalable, cost-effective DNA extraction method. A bead-beating protocol was used to homogenize small amounts of leaf tissue from *Arabidopsis*, camelina, maize, sorghum, soybean, tobacco, and wheat, as well as developing soybean seeds. Samples were processed in 96-well plates using a standard CTAB (cetyltrimethylammonium bromide)-based extraction solution. DNA concentration was measured, and DNA quality was estimated using the A260/A280 and A260/A230 ratios and gel electrophoresis. The extracted DNA was then used for PCR assays to amplify targeted endogenous sequences, detect Cas9-edited sequences, and estimate gene copy numbers. This workflow enabled a single worker to process up to 960 samples per day. The total DNA yield was 2–3 µg for leaf and 0.5-1 µg for soybean seeds with good quality, which was sufficient for multiple PCR reactions. The DNA was successfully used for the intended PCR assays. The extracted DNA was suitable for reliable, downstream PCR-based genotyping. This method supports diverse analyses, including routine amplification, identification of genome edits, and copy number analysis.

## 1. Introduction

PCR-based genotyping is commonly used in plant molecular analyses and is required for many large-scale genetic studies and breeding programs [1,2]. These include quantitative analyses for Quantitative Trait Loci (QTL) using segregating or inbred populations, as well as Genome-Wide Association Studies (GWAS) on natural variation panels. In plant breeding, genomic selection programs often use marker-assisted selection (MAS) on thousands of individuals over 3 to 5 generations. Recent advances in plant genome editing also necessitate large-scale genotyping, with the required numbers of individuals increasing significantly when the desired outcome is infrequent. As an illustration, to produce three T0 plants with a specific editing outcome, it requires producing and genotyping 3–4 (for mutagenesis), 30 (for deletions), 100 (for HDR-mediated integration), or up to 1,000 (for oligonucleotide-mediated editing) T0 plants [3]. The subsequent screening of somatic-edited plants to detect inheritable mutations in the second generation requires genotyping thousands more individuals.

To accommodate the PCR-based genotyping of large populations, a high-throughput DNA extraction method capable of handling many samples is essential. Automated liquid handling platforms (e.g., Biomek 4000, Beckman Coulter, Inc.; MagC 384, 3CR Bioscience) can process a large number of samples effectively but are expensive. Many research laboratories, often with limited labor, time, and budget, need large-scale DNA extraction only on a temporary basis, making investment in this specialized equipment challenging. Therefore, a simple, low-cost DNA extraction method applicable to most molecular laboratories would be more effective for much of the PCR-based genotyping work.

The most common DNA extraction protocols suitable for routine laboratory applications, including enzymatic digestion, adapter ligation, PCR, and sequencing, involve either detergent (CTAB, cetyltrimethylammonium bromide- or SDS, Sodium Dodecyl Sulfate) [4,5] or commercial purification kits (affinity-based). Commercial kits yield high-purity DNA suitable for quantitative analyses (e.g., qPCR or next-generation sequencing), but their per-sample cost is often around US$10 for tube based or around US$2 for plate-based kits. In contrast, CTAB-based protocols are less expensive and provide reliable DNA quality for most qualitative applications, such as routine PCR-based genotyping. The CTAB method has a long history of use for various laboratory applications, including DNA blotting (e.g., Southern blot and RFLP, which require high amounts of DNA) and PCR-based genotyping (which uses small amounts of DNA). Modifications to the standard CTAB protocol, such as the addition of polyvinyl pyrrolidone (PVP), high salt, and 2-mercaptoethanol, help remove polysaccharides and polyphenols and break down proteins. More recently, the CTAB method has been scaled up into a “semi-manual” format with the use of an automated liquid handler, capable of handling 798 samples per day by an average worker, which has proven suitable for genomic DNA sequencing [6].

We present a simpler protocol that requires only common lab instruments (found in most molecular genetics labs) and inexpensive consumables and chemicals. This plate-based CTAB method is simple enough that high school students or other short-trained personnel can extract DNA from 798–960 samples per day. The resulting DNA is reliable for common PCR-based applications, such as routine PCR for endogenous genes, Cas9-edited genotyping, and quantitative PCR (qPCR) for copy number determination. For soybean seeds, only a small amount of tissue is needed, allowing the same seed to be used for both genotyping and downstream molecular assays.

## 2. Materials and Methods

### 2.1 Sample collection

Two subsets of leaf samples were collected from *Arabidopsis* (Columbia), camelina (Suneson), maize (B104), sorghum (TX430), soybean (Thorne), tobacco (*Nicotiana benthamiana*), and wheat (CBO37). The samples were extracted using two methods: a modified plate-based CTAB method and a commercial column-based method (Qiagen DNeasy Plant Mini Kit 69104). For leaf tissue, a 1 cm diameter disk (∼0.8 cm^2^, ∼60–100 mg fresh weight) was sampled with a leaf puncher and kept on ice during collection and transport. Soybean seed samples were collected at the R5.5 (half-size seed), R6 (full-size seed), and R8 (mature seed) developmental stages [7]. To obtain genomic DNA from the embryo only, the seed coat (containing maternal DNA) was removed. A small chip (15–20 mg) from the embryo was collected using a sharp blade.

### 2.2 DNA extraction

The CTAB protocol was modified for plate-based extraction (Figure 1). The reagents are CTAB extraction buffer (containing 2% hexadecyltrimethylammonium bromide - CTAB, 100 mM Tris, 20 mM EDTA, 1.4 M NaCl, and 1% polyvinylpyrrolidone - PVP), chloroform, isopropanol, and 70% ethanol. Approximately 70–100 mg of leaf tissue, or 10-20 mg soybean seed chip was collected into 1.2 ml 12-well tube strips in a rack (VWR 82006-702). The samples were ground in 400 µl of CTAB extraction buffer for 5 minutes at 2500 RPM using a bead beater (Mini-BeadBeater, Biospec Products Inc.), with 4–6 beads for young soybean, tobacco, and *Arabidopsis*, and 7–10 beads for young monocot leaves (Biospec Zirconia Beads, CAT 11079124zx). After collecting the ground tissue into the bottom of the plates via a flash spin, the samples were incubated at 65 °C for 20 minutes. Then, 300µl of chloroform was added to each sample, followed by centrifugation at 4000 RPM (equivalent to 2272 RCF swing rotor with the Sorvall Legend XFR, Thermo Scientific) for 10 minutes in a swinging-bucket centrifuge. Two hundred microliters of the upper phase was transferred to a tube strip pre-filled with 250 µl of isopropanol and mixed well using the same pipette tips to save supplies. The plates were then centrifuged for 20 minutes at 4 °C. The samples were washed with 400 µl 70% ethanol. After overnight drying at room conditions, the samples were resuspended in 200 µl of water for leaf samples or 50 µl for seed samples. The suspended samples were stored in an ice bucket for at least two hours and then mixed well using a multichannel pipette before use in PCR or other applications. DNA extraction using DNeasy Plant Kits (Qiagen) was performed according to the manufacturer’s protocols for a selected set of samples. This set served as a reference for assessing DNA quality and quantity relative to the initial tissue mass.

**Figure 1.**
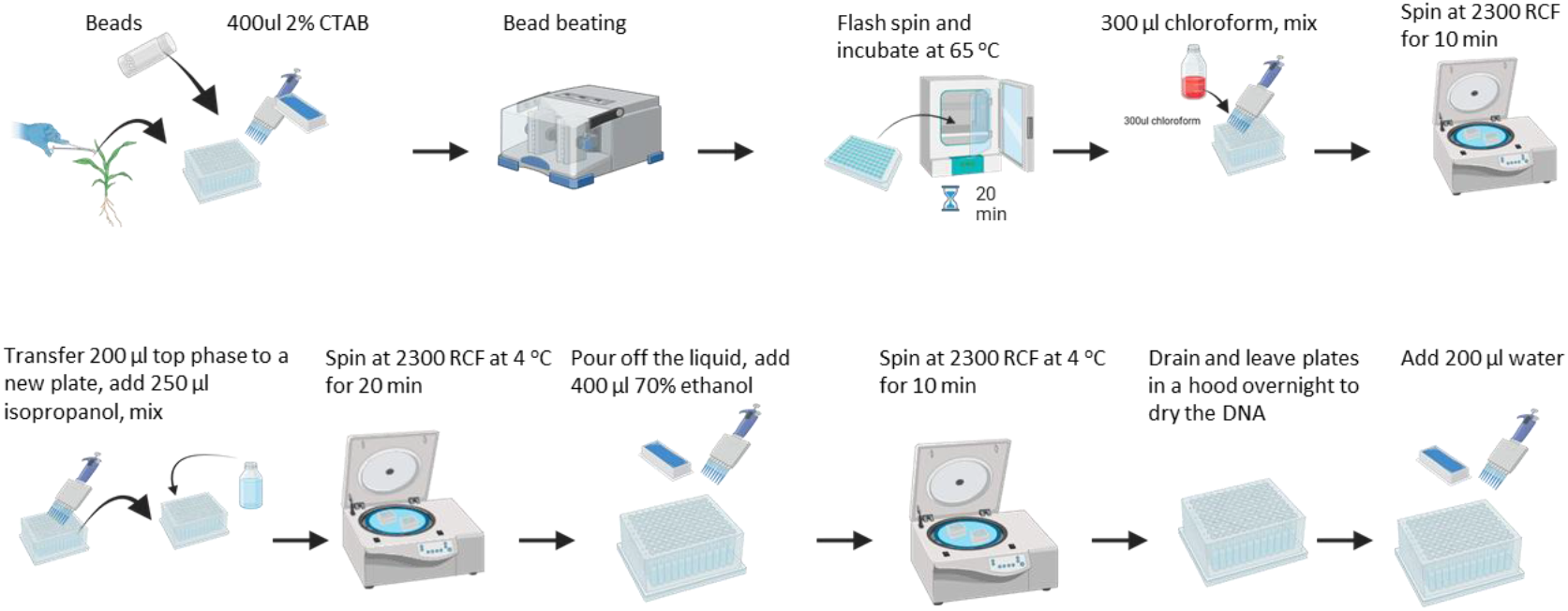
CTAB-based DNA extraction method optimized for multi-plate processing. The extraction process takes 70–80 minutes per two plates and can be conducted in batches to increase efficiency. Faster drying is achieved by inverting the plates during the final draining step. After dissolving the DNA in water, a brief spin removes bubbles to ensure thorough pellet rehydration. Following extraction, the DNA is stored at 4°C for 2 hours and a final DNA solution mixing is needed to improve its resolution before use in a PCR.

DNA Yield and Quality Assay. DNA yield was measured with a spectrophotometer (Nanodrop 2000, Thermo Scientific Inc.), which reported the DNA concentration and the A260/280 and A260/230 ratios. DNA quality was further verified using electrophoresis on a 0.8% agarose gel in TAE running buffer. As the study’s purpose was to evaluate a plate-based method for simplicity, we did not dilute the DNA to a standard concentration for PCR, but instead used 2 µl of template for every sample.

### 2.3 Genomic locus selection, PCR and quantitative PCR

A set of genes from *Arabidopsis, Camelina*, maize, sorghum, soybean, tobacco, and wheat were selected from the literature. These included the reference gene *ATP binding transporter* Glyma.12G020500 [8] for quantitative PCR (qPCR). To enable gene copy number determination, soybean genes were chosen for their presence at multiple genomic locations. Specifically, a set of genes from the GmNFY-A and Linoleate 9-lipoxygenase families, known to exist in 2 and 8 loci, were selected for this purpose (Table 1). For the genotyping of Cas9-edited lines, DNA was extracted from plants previously used in the analysis of a Cas9-derived promoter bashing population [9]. Primers were designed using Primer-3 (https://bioinfo.ut.ee/primer3-0.4.0/) with predefined parameters for GC content, amplicon length, and annealing temperature. The primer sequences were subsequently blasted against species-specific genomic databases (Phytozome for soybean, *Arabidopsis*, sorghum, and maize; Ensembl Plants for tobacco) to identify unique binding sites. Primers were only selected if they were predicted to produce unique amplicons (yielding the same PCR product from different alleles) and had no potential for non-specific binding (Table 1).

**Table 1.**
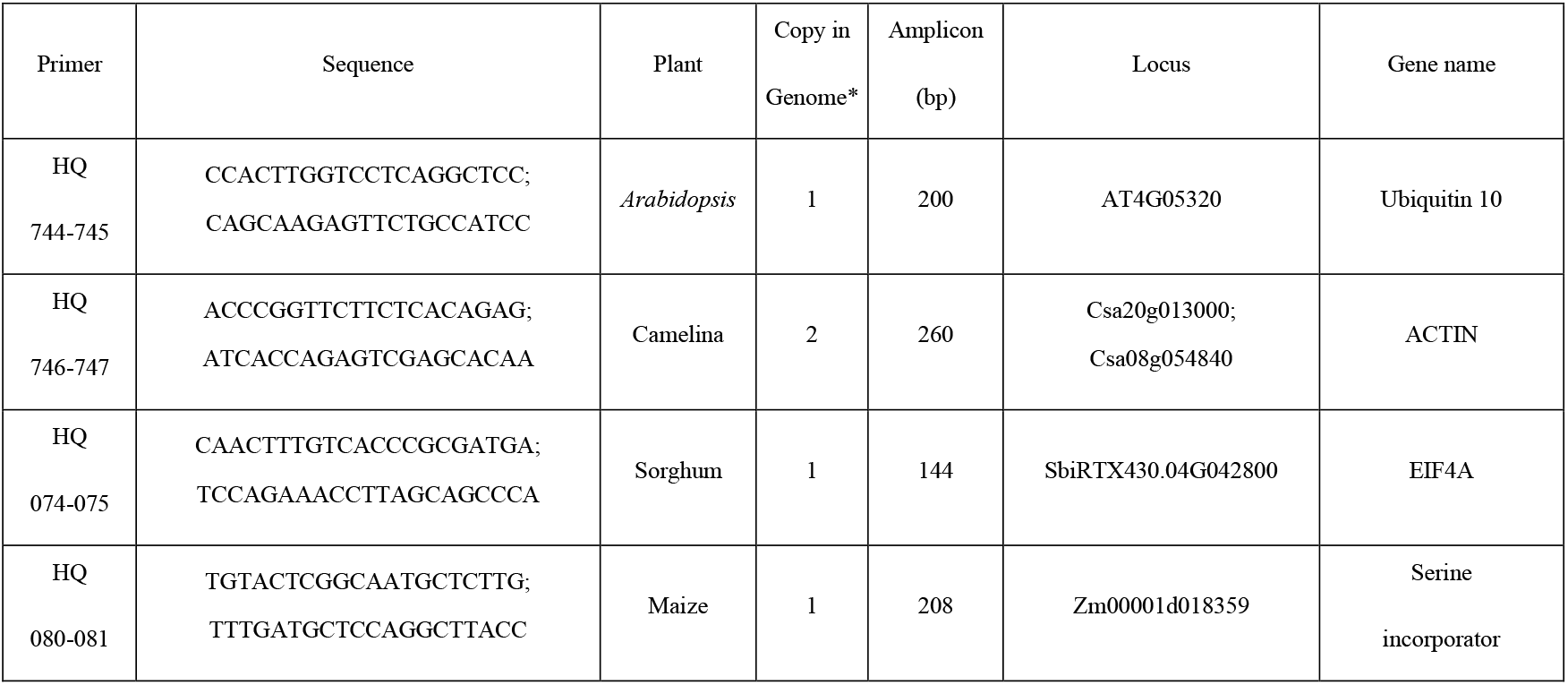
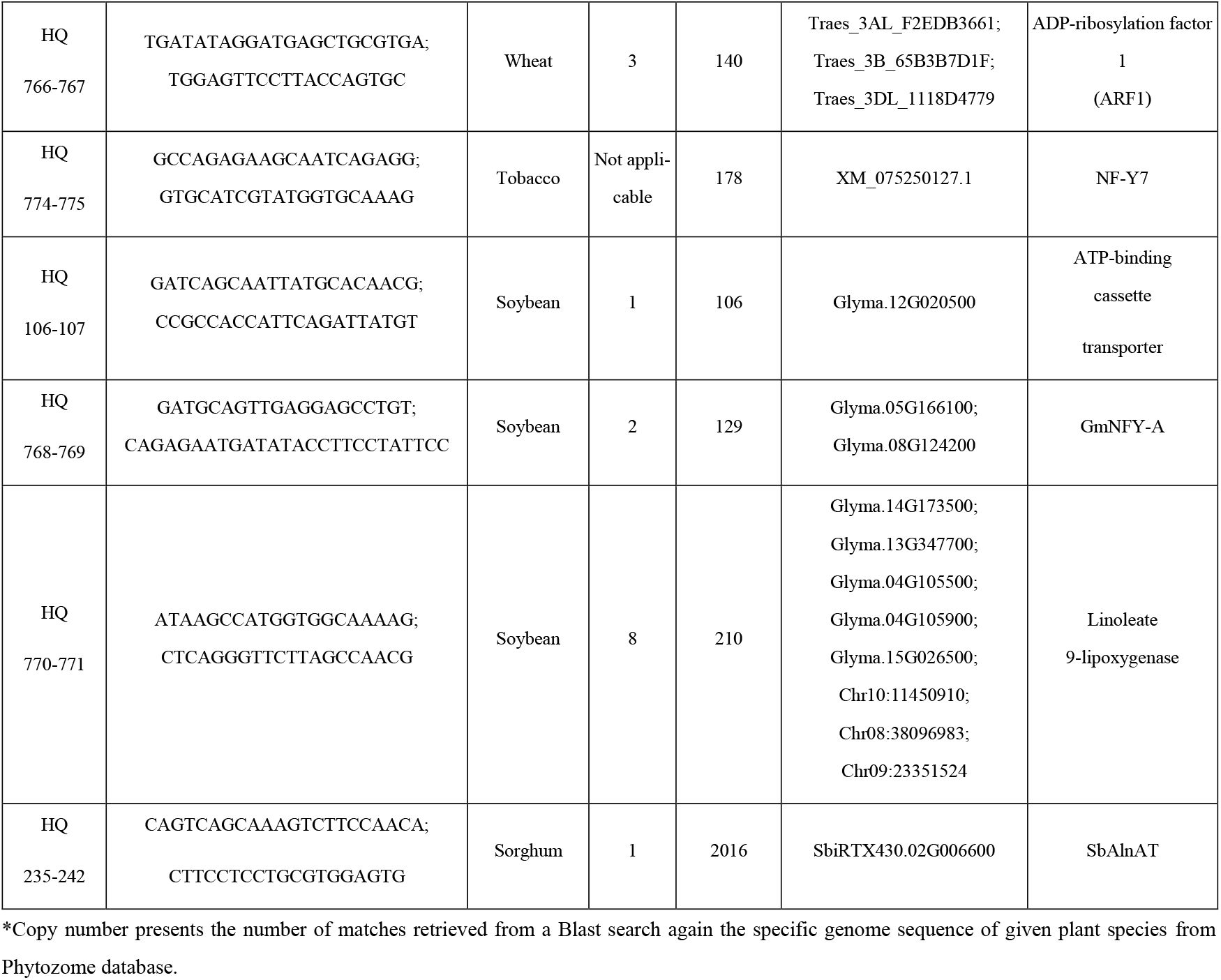
Primers for PCR amplification of endogenous genes and genotyping of genome-edited sorghum lines.

Standard polymerase chain reactions (PCRs) were performed using the GoTaq Master Mix (Promega M7822) in 20 µL reaction volumes, consisting of 10 µL of 2x GoTaq master mix, 2 µL of DNA template, 2 µL of 10x primer mix, and 6 µL of nuclease-free water. The thermal cycling protocol was as follows: (i) an initial denaturation step at 95 °C for 2 min; (ii) 35 amplification cycles of 95 °C for 15 s, 55 °C for 30 s, and 72 °C for 1 min; and (iii) a final extension at 72 °C for 5 min, followed by a holding stage at 4 °C. PCR products were then analyzed by 0.8% agarose gel electrophoresis in TAE buffer. For qPCR, the reaction mixtures were prepared using SYBR™ Green Universal Master Mix (Thermo Scientific 4309155), and reactions were run on a Bio-Rad CFX96 system. Relative gene expression was normalized to the single-copy reference gene Glyma.12G020500 [8]. The gene expression ratio of the target genes to the reference gene was determined for three biological replicates. The copy number was estimated using a chi square (X^2^) test.

## 3. Results

### 3.1. CTAB plate-based DNA extraction method

In the standard tube-based protocol [10], plant samples are ground in liquid nitrogen and then incubated in a CTAB extraction buffer for 30 minutes at 65°C. The DNA is extracted using chloroform, precipitated in isopropanol with high salt, and the DNA pellets are then washed with ethanol. For this protocol, centrifugation is performed at a high speed, often up to 14,000 revolutions per minute (rpm), for 10 minutes. For our present plate-based extraction method, the amount of plant sample is often much smaller (typically 50–100 mg), especially for a seed chip, which can be as small as 10–20 mg. Many lab centrifuges for plates have a maximum speed of only 4,000 rpm for swing rotors, resulting in a relative centrifugal force (RCF) of less than 3,000 g, which is considerably lower than the speed recommended for the tube-based protocol.

We performed a side-by-side comparison using a manual tube-based DNeasy kit from Qiagen and a CTAB plate-based extraction for leaf samples from *Arabidopsis*, camelina, maize, sorghum, soybean, tobacco, and wheat. The quantity of DNA obtained was similar between the methods when using the same amount of tissue (50–100 mg of leaf tissue), yielding a total of about 2–3 µg of DNA. The soybean seed chips (from growth stages R5.5 to R8), however, were relatively small (∼10–15 mg), yielding a lower DNA amount of 0.3-0.5 µg.

The quality of DNA extracted from leaf samples of various common crop species using a plate-based CTAB protocol was evaluated and compared to a column-based kit (Plant DNeasy kit, Qiagen). The plate-based method did not include a RNase treatment, as RNA contamination does not interfere with standard PCR applications. Traces of RNA were visible in the samples from the agarose gel electrophoresis (Figure 2), indicating a slight difference in purity.

**Figure 2.**
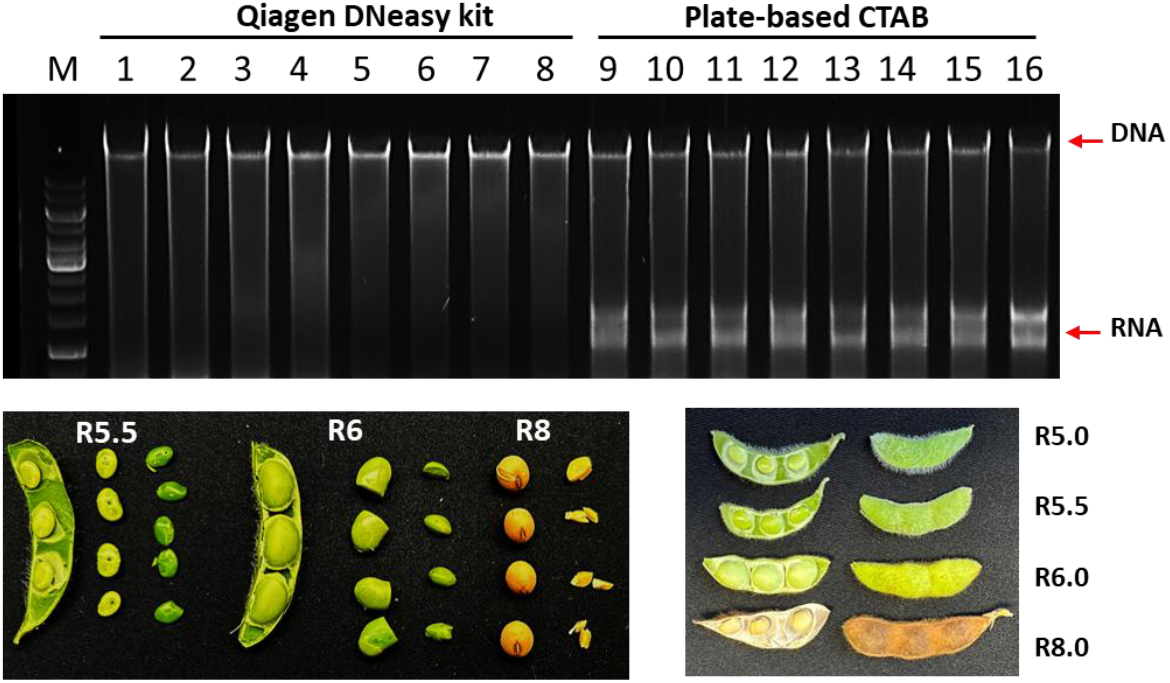
Agarose gel electrophoresis of total DNA extracted from various plant species. Total genomic DNA was extracted from the leaves of seven plant species and from soybean seed using a CTAB-based method. Fifty nanograms of DNA was loaded into each well for electrophoresis. Lanes 1-7 and 9-15 correspond to leaf DNA from *Arabidopsis*, camelina, maize, sorghum, soybean, tobacco, and wheat. Lanes 8 and 16 represent DNA extracted from soybean seeds at the R6 developmental stage. The bottom panel of the figure illustrates the key soybean seed development stages of R5.5, R6 and R8, which show the full pod, the harvested seeds, and the seeds without seed coat to be used for DNA extraction.

The DNA from both methods showed comparable quality and quantity (Table 2 and Figure 2). Electrophoresis revealed a single, distinct high-molecular-weight band with minimal streaking, indicating minimal shearing in both extraction methods. The plate-based method yielded good quality DNA with little to no smear, which is confirmed by the electrophoretic results. Spectrophotometric analysis using a Nanodrop revealed the characteristic absorption readings at 280, 260, and 230 nm, which were used to calculate the A260/A280 and A260/A230 ratios. The DNA purity, as reflected by these indices, may be lower in the CTAB samples due to co-precipitation of other substances during extraction. For assays that require very high DNA purity, such as next-generation sequencing, column-based or other affinity-based purification methods are recommended.

**Table 2.**
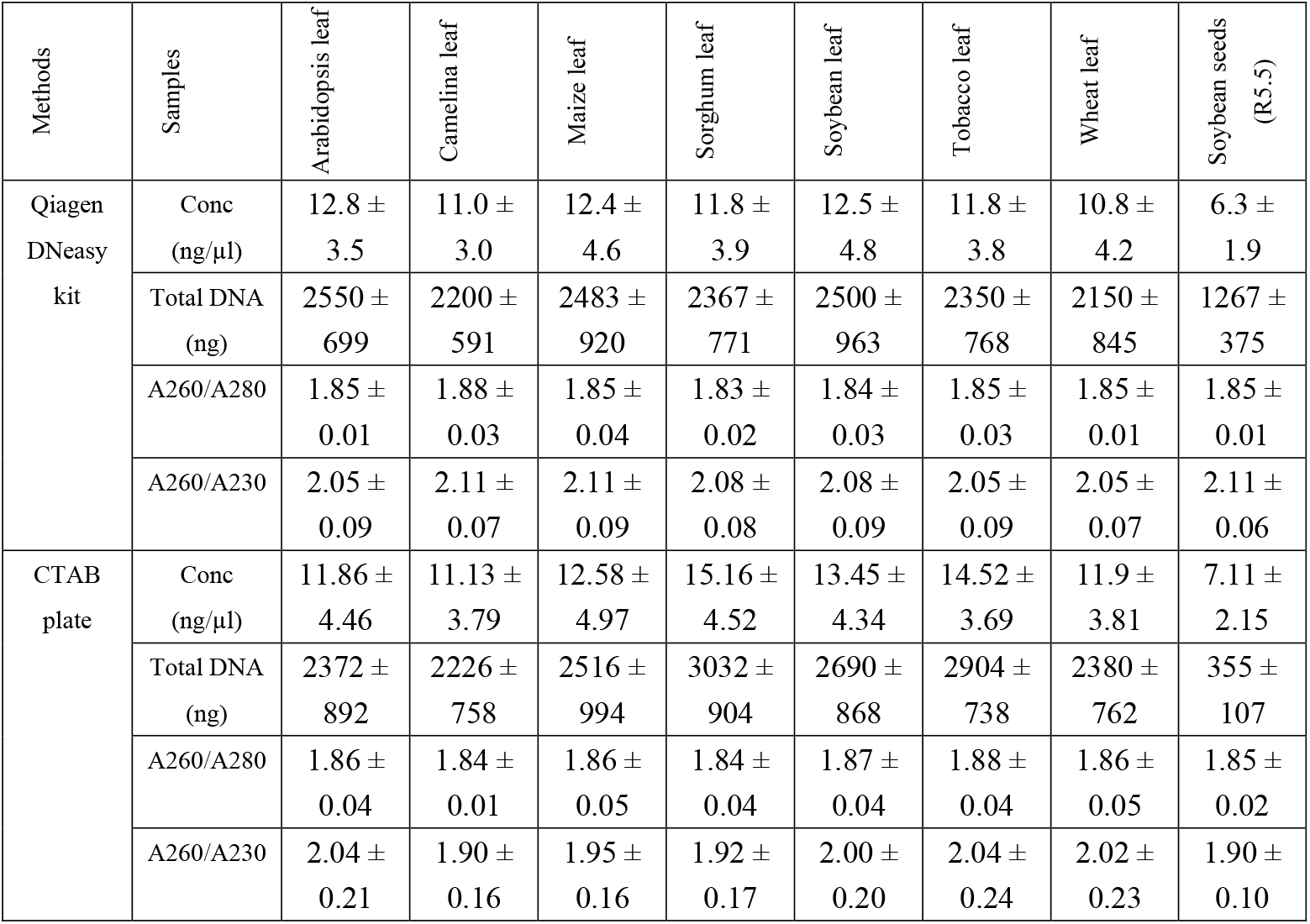
Estimation of DNA quantity and purity using a plate-based extraction method for various crop tissues. DNA from leaf samples (60–100 mg fresh weight; ∼0.8 cm^2^) were resolved in 200 µl of water and DNA from soybean seed chips (15–20 mg), collected at growth stage R5.5, were resolved in 50 µl of water. Twelve samples of each tissue type were extracted using a 96-well plate. Data are presented as the mean ± standard deviation (*n*=12).

The total DNA yield from the plate-based method ranged from 2 to 3 µg for leaf, with concentrations between 10 and 15 ng/µl (Table 2). This amount is sufficient for most standard PCR-based molecular assays. In a given assay that requires higher concentrations, the DNA can simply be eluted in a smaller volume of water. We did not attempt to quantify the DNA concentration of the seed samples at R8 using a NanoDrop spectrophotometer. Measurements at this stage would not have been reliable due to the high coprecipitation of polysaccharides, which can produce erroneously high readings. For seed genotyping, a small portion of the seed was used for DNA extraction, while the remainder could be used for regenerating plants or for other phenotypic assays. Since the seed coat has the genotype of the mother plant, it was removed prior to DNA extraction (Figure 2) to avoid contamination from maternal DNA. Because the amount of seed tissue is small (5–15 mg), it yielded a significantly lower amount of DNA compared to leaf tissue.

### 3.2. Use of DNA for standard PCR

For routine PCR screening, we use GoTaq Green Master Mix (Promega M7823), which is available in a high-volume format sufficient for 1,000 standard 50 µl reactions. Given our needs, we typically perform 20 µl reactions, allowing for up to 2,500 reactions per bottle. This reduced reaction volume did not cause any noticeable decrease in PCR product yield, which was confirmed by clear band visualization on a gel for different plant species (Figure 3A).

**Figure 3.**
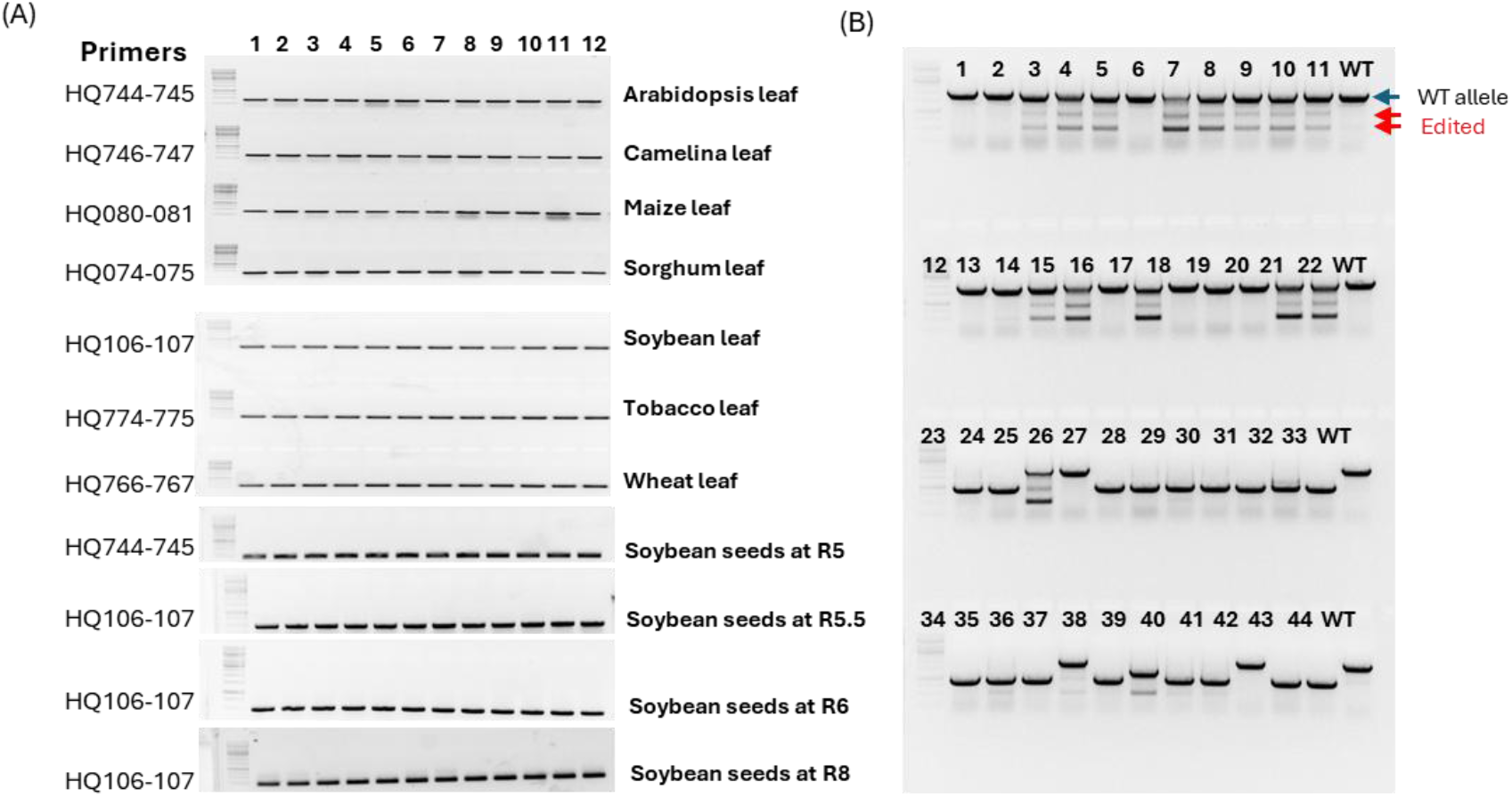
Application of plate-based DNA in PCR assays. The figure demonstrates the versatility of plate-based DNA across four different PCR assays. (A) Representative standard PCR results for DNA from various plant species, using primers listed in Table 1. The numbers shown on top represent the 12 individual samples in the same set of tissues for consistency. (B) Results from PCR genotyping of 44 T1 plants derived from the Cas9-edited sorghum event TZ152-2B [9]. A 2 kb PCR product (primers HQ235-242), amplified from the wild-type (SbAlaAT) promoter, is visible in the wild-type (WT) control lanes. Genotype results for the edited lines were categorized as non-edited (lanes 1, 2, 6, 12, 13, 16, 18, 19, 20, 26, 42) showing a single WT band, somatically edited (lanes 3, 4, 5, 7–10, 14, 15, 17, 21, 22, 25, 37, 39) showing multiple bands, and germline-edited (lanes 23, 24, 27–36, 38, 40, 41, 43, 44) showing distinct mutant bands.

When performing high-throughput PCR, DNA template concentration was not standardized prior to loading. Instead, we used a consistent volume of 2 µl of template DNA for all samples, including those from smaller tissue types such as seed chips. The DNA extracted using our plate method is sufficient for 100-200 such reactions per which about 10-20 ng of DNA template is used. For additional reactions, the DNA can be further diluted without affecting gel visualization quality.

Seed chipping proved effective for genotyping while preserving the seeds for planting or other biochemical assays. For this method, a small tissue amount (10-20 mg) was used, which yielded a small quantity of extracted DNA. In our PCR assays, we did not normalize the DNA input but used 2 µl of the template, corresponding to approximately 5-10 ng of DNA. The total PCR volume was 20 µl, with half of the product loaded onto a gel for analysis. The protocol yielded clear and consistent PCR bands across all 12 tissue samples tested for both leaf tissue from different plant species and soybean seed developmental stages

Plant genome editing has been used extensively for functional genomics, from validating candidate genes identified through high-throughput genetic analyses to improving crops. Since inheritable mutations occur at a low frequency, a large population of edited lines must be screened to identify both somatic and germline mutations. This screening process requires generating and genotyping a large population. Using a recent sorghum materials [9], we performed PCR on DNA extracted from lines which had been edited for the gene SbAlaAT. The results, shown in Figure 3B, demonstrate clear mutation patterns, with edited genome showed somatic and germline deletions. The consistent and clear bands across all samples indicate the reliability of our extracted DNA. For this study, we used a simplified DNA extraction protocol consisting of a CTAB extraction, chloroform purification, isopropanol precipitation, and ethanol washing. We omitted the RNAse treatment step to make the procedure faster and more cost-effective and observed no adverse effect on our PCR results

### 3.3. Copy number determination in soybean

Gene copy number is a critical area of research in genomics, evolution, and genetic engineering. For example, in soybean, copy number variations in the *rhg*1 locus are correlated with nematode resistance [11]. In transgenic plants, higher copy numbers can be associated with gene silencing and epigenetic effects, while varying copy numbers can have either a positive or negative effect on expression [12,13]. For genetic engineering applications, determining the copy number of transgenes can significantly reduce the amount of downstream genetic screening and increase the functional stability of the transgenes in transgenic plants. Consequently, maintaining a low, and ideally single, copy number is a primary objective in genetic engineering for developing stable and homozygous traits, particularly in gene stacking [3]. Copy number can be determined using PCR based DNA sequencing and qPCR [14,15].

Here, we explored the use of a plate-based CTAB extraction method combined with qPCR to determine gene copy number. We used endogenous genes with known copy numbers of 1, 2, and 8 in the soybean genome (Table 1). Results from the qPCR were normalized using a single-copy reference gene [8]. Figure 4 show a clear correlation between the gene copy number and the normalized qPCR signal, with signal ratios that correspond well to the predicted copy numbers.

**Figure 4.**
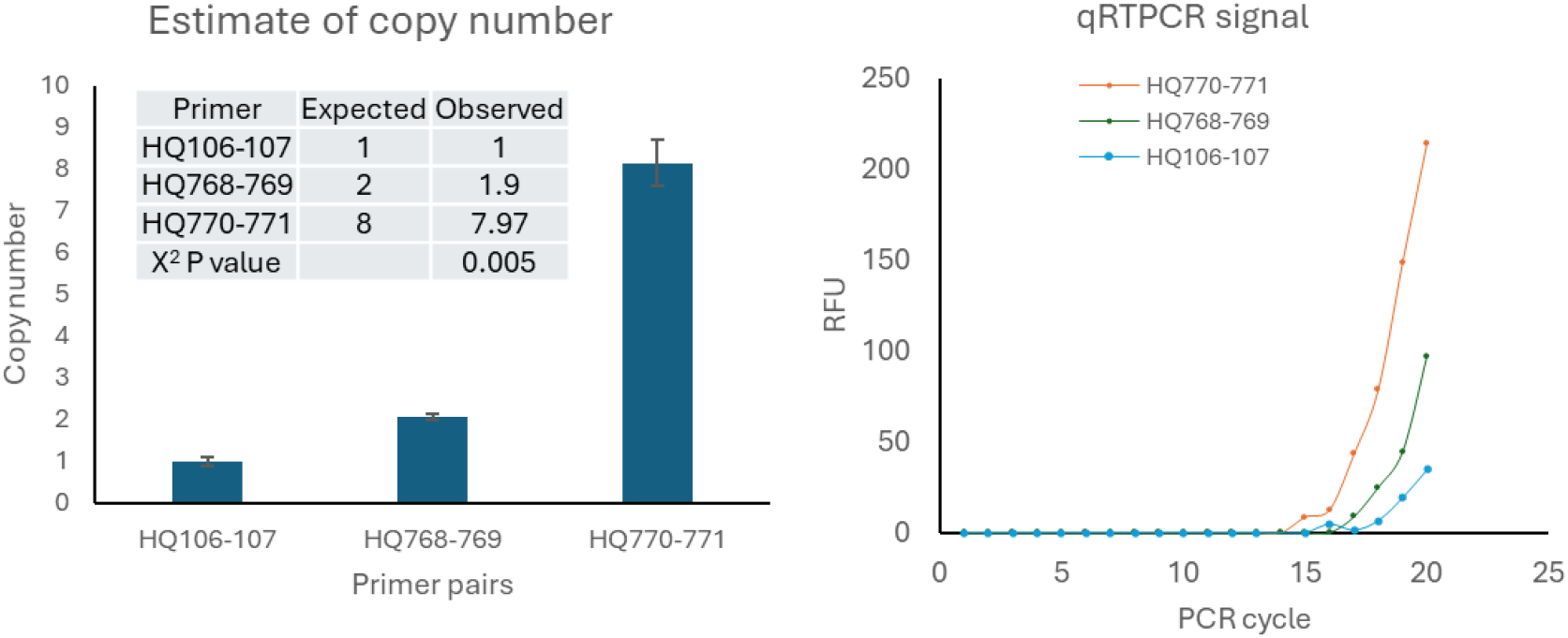
CTAB plate based extracted DNA was used for determination of gene copy number in soybean. The expected copy number was retrieved from soybean genome database hosted in Phytozome. qPCR was used to quantify the copy number and the observed values for each gene were normalized to the single copy number ATP-binding cassette transporter. The insert table in the left figure shows the X^2^ test with high confidence level (n=3).

### 3.4. Efficiency, time and financial cost

As the protocol is relatively simple and use of low cost chemicals and consumables, the major proportion of the total cost is labor cost. In our protocol, a worker could demonstrate 8-10 plates per eight-hour workday. The DNA plates are dried overnight, and water can be added the next day and ready for PCR after two hours and can be stored at -20 °C for a year for later use. The tube strips and transfer pipette tips are universal from commercial suppliers and often cheaper than those purchased from a special kit provider for special equipment such as the ones bundled with a specific liquid handler. This is an advantage over the use of the previously reported RoboCTAB method [6] which use the consumables specifically provided with the automated liquid handling OT-2 system (Opentrons Inc.).

The required equipment is a Tissuelyser (or a bead beater), a centrifuge with plate swing rotor that can spin at ∼2300 RCF, and two multi-channel pipettors (200 and 1000 µl type). The cost per sample was calculated as the labor cost ($US0.14), cost of consumables ($0.14) and the cost of reagents ($0.07). Considering these factors, the normalized cost per sample was $0.35 for the year 2025. This cost is relatively less than the cost estimated recently reported [6] within the same year and reagents, but using the semi-automated protocol using a liquid handler. The liquid handler, however, can increase initial cost significantly and not an investment if the demand is seasonal and periodically in a small lab. The other cost associated with the liquid handler is the consumables that might be expensive and not available in many international markets.

### 3.5. Routine use of the methods in genotyping works

Our method has been used by research laboratories to genotype tens of thousands of samples each year (Table 3). Examples of this work across different crop species including soybean, sorghum, and wheat. The results have been validated in the literature, demonstrating reliable performance across a variety of tissue types.

**Table 3.**
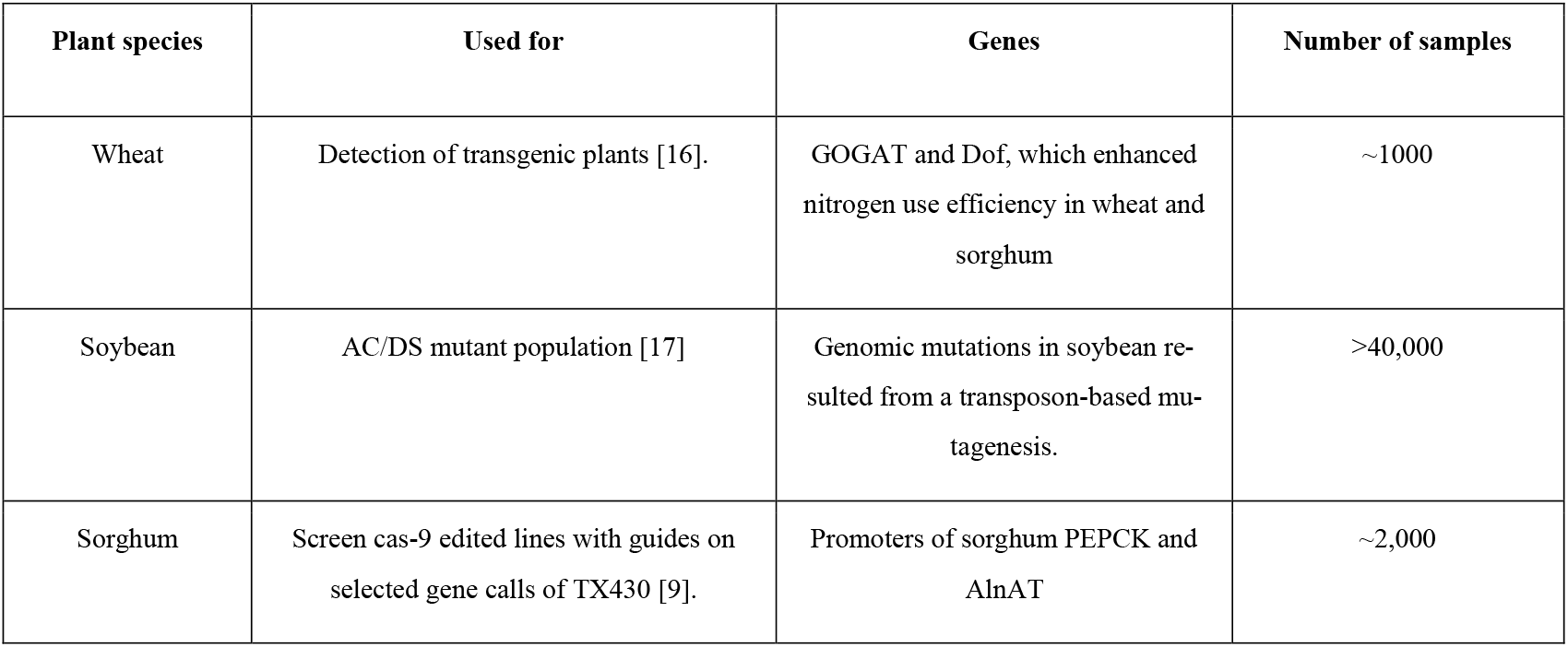
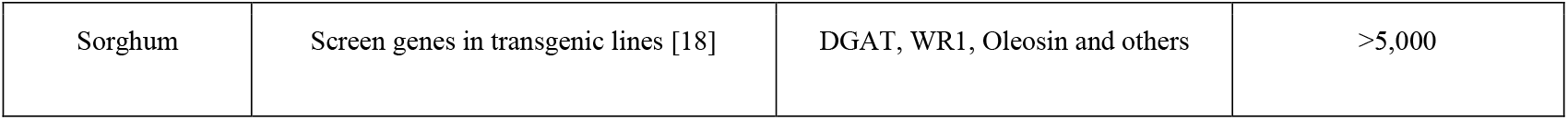
Examples of genotyping works using the CTAB plate-based DNA extraction method for various PCR based assays.

## 4. Discussion

Modified DNA extraction protocols, such as those using CTAB [4] and SDS [5], became standard methods for a wide range of applications, including both PCR-based and hybridization-based genotyping. While these conventional methods yield high-quality and high-quantity DNA, they are often time-consuming and labor-intensive, especially when processing large numbers of samples. For studies involving large populations—such as mapping, genome editing, or genomic selection populations—high-throughput DNA extraction is necessary. To meet this demand, automated DNA extraction instruments have proven effective (Boucher St-Amour et al., 2025). These systems most often employ affinity-based methods, which produce the high-purity DNA required for sensitive molecular assays. Such instruments are available from major providers including Thermo Fisher Scientific and Qiagen. Although they require a significant initial investment, automated platforms are a cost-effective solution for specialized, high-demand DNA services.

To simplify the preparation of DNA for routine use in many research laboratories, a recent report by Boucher St-Amour et al. (2025) describes a method that combines a modified CTAB extraction with a liquid handling system. This technique is particularly suitable for labs processing smaller populations, with annual volumes ranging from several hundred to tens of thousands of samples. The protocol employs a semi-manual workflow, where an operator prepares the samples and manages the mini-beater and centrifuge, while the liquid handling system automates the liquid transfer steps. This approach is not only cost-effective but also allows an average worker to process 768 samples (8 plates) daily, yielding high-quality DNA for sequencing applications. Given its advantages, this method is a suitable option for any facility equipped with a liquid handling system.

In this report, we describe a simple, plate-based CTAB protocol that can be applied to various research and development projects requiring a large number of DNA samples. The method yielded 2–3 µg of DNA from soybean *Arabidopsis*, camelina, maize, sorghum, soybean, tobacco, and wheat leaves and approximately 0.3-1.5 µg of DNA from soybean seeds. The obtained DNA is suitable for PCR analyses for routine gene detection, identification of genomic mutations generated by genome-editing tools, and determination of the copy number of endogenous genes or transgenes in the plant genome.

Our method is a modified CTAB protocol that does not require an automated liquid handling system, which is unavaiable in most laboratories. It also does not require the use of RNase or mercaptoethanol during the extraction process or DNA quantification before PCR. The protocol works well for soybean chips, making it suitable for anyone wanting to couple PCR with additional seed composition assays or for saving seeds for later sowing.

Cost savings are realized through the higher volume of samples a worker can proceed a day, the elimination of RNase and mercaptoethanol, and the absence of a liquid handling system. Compared to tube-based DNA extraction, this method also offers a higher volume of sample handling, a significant reduction in labeling, and a lower chance of errors from labeling and liquid transfers.

We have not tested the DNA for sequencing, but since the protocol is relatively similar to the roboCTAB method, the quality of the DNA is expected to be good for genotyping by sequencing, as previously reported [6]. However, because the protocol does not use a high-salt DNA precipitation step, we expect some contamination from plant tissues. For other assays requiring higher purity DNA, we do not recommend the current protocol.

